# Duration discrimination in the bumblebee *Bombus terrestris*

**DOI:** 10.1101/2025.02.03.636111

**Authors:** Alexander Davidson, Ishani Nanda, Anita Ong Lay Mun, Lars Chittka, Elisabetta Versace

**Affiliations:** School of Biological and Behavioural Sciences, Department of Biological and Experimental Psychology, Queen Mary University of London, London (UK); Nanyang Technological University, Singapore

**Keywords:** Time perception, duration discrimination, interval discrimination, bees, bumblebees

## Abstract

The ability to process temporal information is crucial for animal activities like foraging, mating, and predator avoidance. While circadian rhythms have been extensively studied, there is limited knowledge regarding how insects process durations in the range of seconds and sub-seconds. This study aimed to assess bumblebees’ (*Bombus terrestris*) ability to differentiate the durations of flashing lights, and use this information in a free-foraging task. Bees were trained to associate either the long or short-duration stimulus with a sugar reward versus an unpalatable solution until reaching a criterion, and then tested without sucrose solution with the same stimuli. In Experiment 1, we tested the ability to discriminate between a long stimulus (2.5 or 5 seconds) vs a short stimulus (0.5 or 1 second). The bees learned to discriminate between the two stimuli. To check whether bees solve the task without using the absolute difference in proximal stimulation as a cue, we ran a second experiment. In Experiment 2, the flashing stimuli were presented for the same total amount of time in a cycle. Bees could discriminate between durations even when the overall amount of stimulation in each presentation cycle was the same. This shows general learning abilities in bumblebees, that can discriminate seconds/subseconds intervals in visual flashing stimuli. This reveals an insect’s ability to use non-naturalistic stimuli and temporal cues in free foraging.

## 1. Introduction

Animals can use temporal cues such as the duration of time intervals to drive their choices. For instance, hummingbirds and bees time flower visits according to the rate as which specific flowers replenish their nectar supply [1–4]. Effective communication can also require temporal encoding and decoding of intervals, such as in the waggle dance of bees [5–7], or the mating calls of crickets, where different durations correspond to different distances and species [8]. Time scales of intervals relevant to animals vary by orders of magnitude, from years to fractions of a second. The mechanisms behind time processing in different time scales may be manifestations of a generalised ability to keep time [9], or alternatively, there may be dedicated mechanisms for specific time scales [10]. Longer time scales such as circadian cycles depend on protein synthesis and degradation [11] that cannot account for shorter intervals in the scale of sub-seconds or seconds, which are often relevant for foraging and communication. Here we focus on the ability of an insect, the bumblebee *Bombus terrestris*, to discriminate between visual time intervals in the scale of sub-seconds (0.5 s) to seconds (2.5-5 s) in a free-foraging task. Bees’ ability to discriminate between different timings of flashing stimuli might suggest an adaptability of the bee’s visual system to process general regularities, and their ability to learn using arbitrary temporal cues as relevant information to drive behavioural choices.

One of the first studies looking into the cognition of interval duration was published in the 19th century and investigated the estimation of intervals in humans [12]. An early study with Rhesus monkeys (*Macaca mulatta*) showed the ability to discriminate between two interval durations (1.5 s vs 4.5 s) [13]. Pigeons can use interval timing as a cue for foraging [14]. Rats can also discriminate between durations and track the duration of intervals [15,16]. Little is known on insects. Wasps have the ability to associate a duration (5 vs 30 minutes) of time with a reward [17]. Bumblebees (*Bombus impatiens*) can learn to expect a reward after a given interval (6 or 36 s) [18], although this result was challenged by a work that did not find evidence of temporal control in honeybees (*Apis mellifera*) [19]. Bees are known to be good at visual learning, and have keen visual acuity [20–22] and can be conditioned to reach for sucrose solution with continuous or intermittent reinforcement [23]. A small study conducted on honeybees suggests they are able to discriminate between flashing lights [24]. This makes bees a good model for investigating time processing at smaller time scales using visual stimuli, though this area of research remains largely unexplored.

The present study focuses on duration discrimination of visual stimuli in the sub-second to seconds range, using a classical conditioning paradigm in free moving bees. The sub-second to seconds time scale is relevant for everyday tasks, such as navigation foraging and communication [7,25,26]. We chose these durations based on the visual working memory of honeybees which show a robust working memory of up to 5 seconds [27–30].

## 2. Methods

### (a) Subjects

We obtained bumblebees (*Bombus terrestris*) from Agralan Ltd. Upon arrival, bee nests were transferred into a wooden nest box (Fig. 1). We reared bees at 22–23°C, with at 12 hour light dark cycle. Bees fed on a 20% sucrose solution available ad libitum, and we provided pollen every other day. We tested 41 bees from 10 different colonies with a fully counterbalanced design regarding the rewarded stimulus. In Experiment one we tested 20 bees. We trained 10 were trained to associate a sugar reward with the long duration stimulus and 10 were trained to associate a sugar reward with the short duration stimulus. In Experiment 2 we tested 21 bees. We trained 10 bees on the long duration stimulus and 11 on the short duration stimulus.

**Figure 1.**
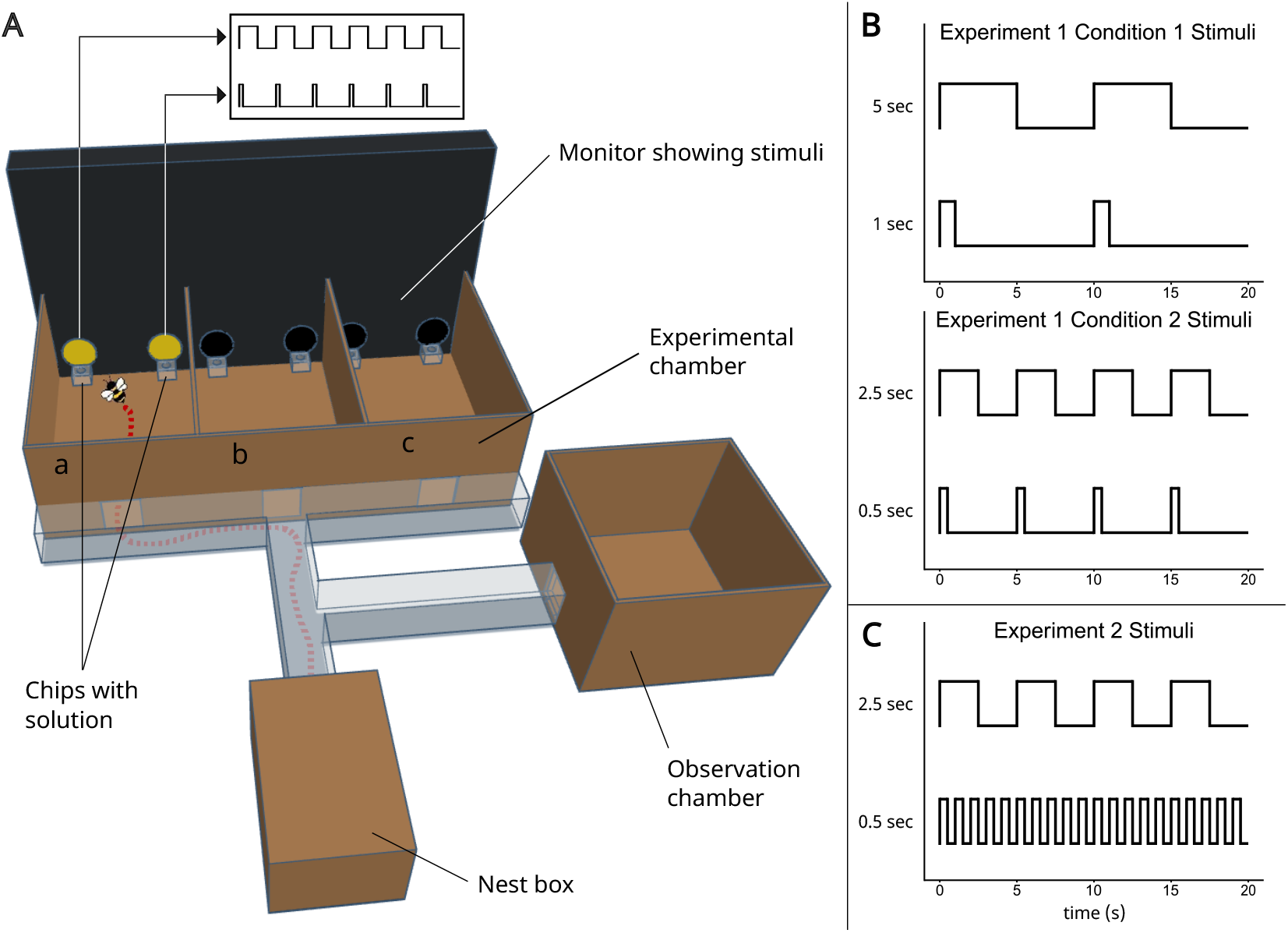
(A) Experimental apparatus. A bee has entered the left compartment (a) of the Experimental chamber. Its route is traced from the Hive box. The bee will then visit compartments b and c, before returning to the nest box. A simplified representation of the stimuli is illustrated above the monitor and shown in more detail in the panels on the right. (**B**) The durations of stimuli in Experiment 1 (5 vs 1 seconds in the top panel, 2.5 vs 0.5 seconds in the bottom panel. The peaks represent the “on” state, where the stimulus is displayed. (**C**) The durations of stimuli in Experiment 2. Over each full cycle of the long stimulus, the sum of “on” states for both stimuli is equal.

### (b) Apparatus

The apparatus (Fig. 1A) included a Hive box, an Observation chamber and an Experimental chamber connected by clear acrylic tunnels with removeable doors to control the movement of bees. Each colony was raised in the Hive box. Bees had free access to the observation chamber, where foragers were identified while visiting a 20% sucrose solution station. The observation chamber (30×30×21 cm) was covered with a clear acrylic sheet so bees could be observed feeding. A single forager per day was allowed into the experimental chamber, where bees were trained in a double choice discrimination task until reaching a criterion and then tested.

The experimental included three identical compartments (a-c in Fig. 1A) (25x17x10.5 cm). The three chambers allowed us to run three independent trials for each foraging bout. An LED strip located above the experimental chamber provided light. Acrylic chips held feeding solutions during trials. We used a computer monitor (ASUS XG258Q, 240 Hz) placed opposite to the experimental chamber to display the stimuli.

### (c) Stimuli and rewards

Stimuli were 4 cm Ø circles in yellow (hex FFFF00) on a black background. Each circle blinked on and off at the given frequency for each experiment. During the training, each stimulus could be used as predictor of the presence of palatable unpalatable food. Pairs of stimuli were used, shown (11.5 cm) apart so that two stimuli were present in each compartment of the experimental chamber (Fig. 1). The feeding chips were filled with 20 µl of 50% sucrose solution or 0.12% quinine solution and paired with the stimuli according to the experimental schedule.

In Experiment 1 we tested bees’ ability to discriminate between different durations of presentation of visual stimuli. We used two conditions (Fig. 1B). In condition 1, we used a 10-second cycle in which the long duration was presented for 5 seconds and the short duration for 1 second, with a simultaneous onset. In condition 2 we used a five second cycle in which the long duration stimulus was presented for 2.5 seconds and the short for 0.5 seconds with a simultaneous onset.

In Experiment 2 we tested bees’ ability to discriminate between different interval durations when the amount of stimulation per cycle was matched for both stimuli (Fig. 1C). We used a 5-second cycle in which the long duration stimulus flashed on and off in 2.5 second intervals, and the short duration stimulus flashed on and off at 0.5 second intervals. This amounted to both stimuli being in the on state for 2.5 of the 5 second cycles.

### (d) Training

Training consisted of repeated foraging bouts to the experimental chamber. A tagged forager was allowed to enter the three compartments (Fig. 1A) consecutively, each compartment taken as an independent trial. In the compartment were two feeding chips – one with 20 µl of 50% sucrose solution and the other with 20 µl 0.12% quinine solution placed immediately in front of the rewarded and punished stimuli respectively. At the start of each trial, the bee was held in the tunnel at the entrance to the compartment for 10 seconds, while the stimuli were visible through the transparent tunnel. The bee was then allowed to enter the compartment, and its choice of stimuli was recorded as the first feeding chip it touched with any part of its body. Once the bee left the compartment, the same procedure was followed for the next two compartments. The right-left stimulus position was counterbalanced between trials. Once the bee left the last compartment, it was allowed back to the nest box. The experimental chamber was cleaned with 70% ethanol solution and paper towel, the chips removed and replaced for the next trials. When the bee came back out of the nest box, the procedure was repeated. Training continued until the bee reached a criterion of 15 correct choices out of the last 20. This corresponds to a p-value = 0.041 in a binomial test.

If a bee showed a side bias (two consecutive incorrect choices on the same side), correction trials were run, which did not contribute to the criterion of success. In correction trials, the rewarded stimulus was presented on the side opposite that of the bias.

### (e) Test

We tested bees in 15 trials using water in both feeding chips (in extinction). To ensure that bees did not stop responding [23], test trials were interspersed with training trials with sucrose and quinine. The compartment used for the test trial was randomized between bouts.

### (e) Data analysis

To assess the discrimination between the stimuli, for each experiment we analysed the population and individual performance. Alpha was set to 0.05.

For the population analysis, we ran an ANOVA to test whether there was a difference in performance between the two conditions of Experiment 1 (long cycle of 10 s vs short cycle of 5 s). Since there was no difference between conditions (F (1,18) = 0.96, p = 0.339; M_10s_ = 0.667, SEM_10s_ = 0.067; M_5s_= 0.6, SEM_5s_ = 0.042; see Figure 2A), we collapsed all the data and used a one sample t-test to analyse bees’ choice of stimuli as a group.

**Figure 2.**
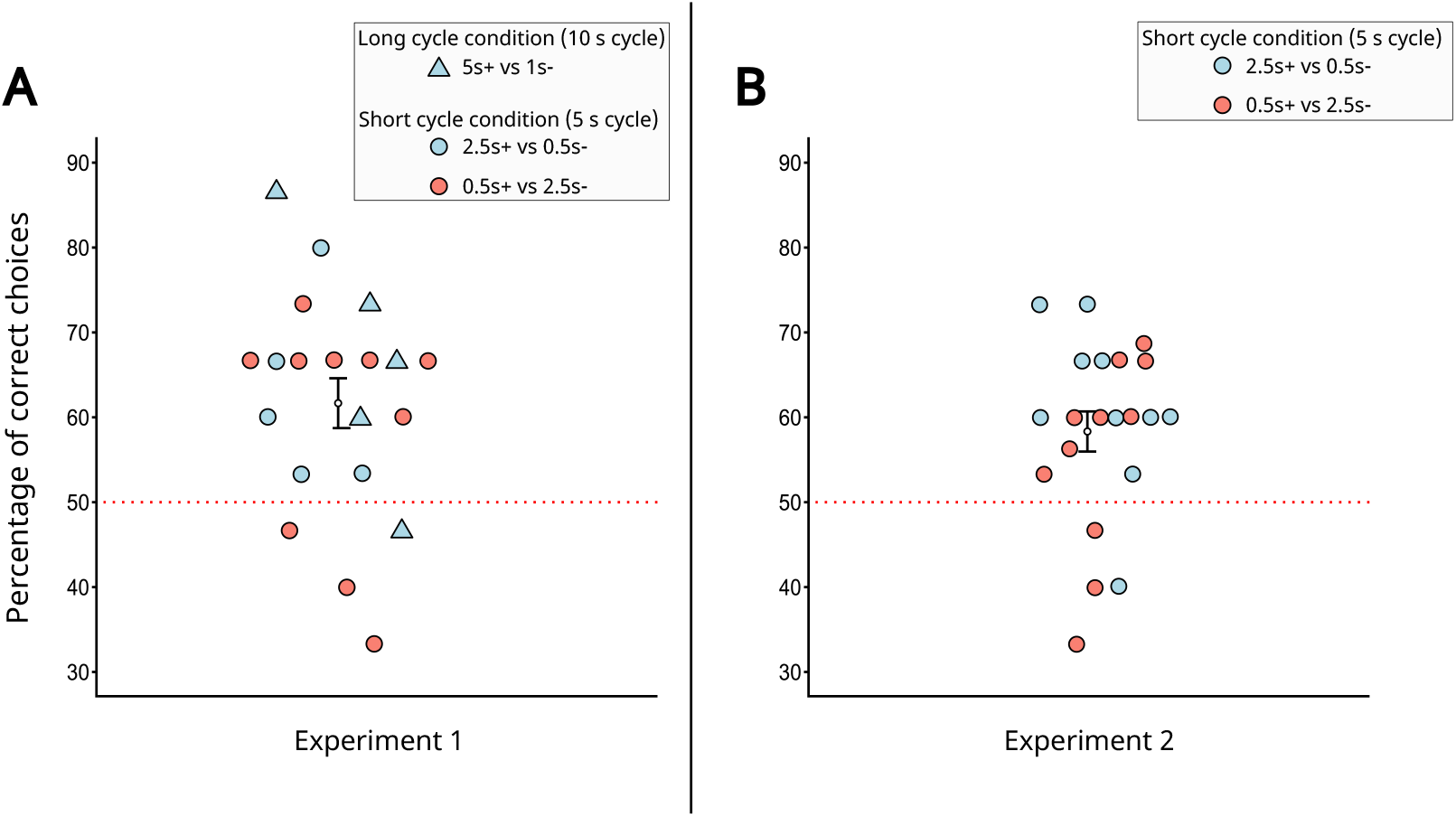
Results of test trials. Each point represents an individual bee. The dashed red line represents the random choice level. (**A**) The percentage of correct choices for each bee in Experiment 1. (**B**) The percentage of correct choices for each bee in Experiment 2.

We used a paired t-test to confirm the absence of any difference between the test trials and the reinforcement trials. We used a paired t-test to check for any preference for the left or right side of the compartments of the experimental chamber. We used an unpaired t-test to check whether there was any different in performance between Experiment 1 and Experiment 2.

To assess individual bees’ performance, we used a binomial exact test with a hypothesised proportion of 0.5 correct responses.

## 3. Results

### Experiment 1

At the group level, bees learned to discriminate between different durations of presentation of visual stimuli (t(19) = 3.972, p < 0.001) (Fig. 2). The mean score for the long + group was 0.647 (SEM = 0.152), the mean for the short + group was 0.587 (SEM = 0.156). There was no difference in performance between the two conditions in experiment 1 (F (1,18) = 0.96, p = 0.339), and no preference for the left or right side of the compartments (t(19) = -1.254, p = 0.225). We found no difference in performance between test trials and reinforced trials during the test phase (t(19) = -0.232, p = 0.819).

At the individual level, 2 out of 20 bees performed significantly above chance level (p_bee 12_ = 0.007, p_bee 15_ = 0.035). Overall, 16 out of 20 bees chose the correct stimulus more than 50% of the cases.

### Experiment 2

At the group level, bees learned to discriminate between the 2.5 second and 0.5 second stimuli (t(20) = 3.528, p = 0.002). The mean score for the long + group was 0.613 (SEM = 0.155, n = 10). The mean score for the short + group was 0.557 (SEM = 0.15, n = 11). There was no preference for the left or right side of the compartments (t(20) = -1.460, p = 0.159).

We found no difference in performance between Experiment 1 and Experiment 2 (t(39) = 1.212, p = 0.233). We found no difference in performance between test trials and reindorced trials during the test phase (t(20) = -0.24432, p = 0.809). There was no difference in performance between the two Experiment 1 and Experiment 2 (t(36.873) = 0.884, p = 0.382).

At the individual level, no bees performed significantly above chance level. Overall,17 out of 21 bees chose the correct stimulus more than 50% of the cases.

## Discussion

Our results show that bumblebees can learn to discriminate between different durations of visual stimuli presented as flashing lights to guide their foraging choices. We tested bees using flashing yellow circles presented for different durations, paired with appetitive and aversive feeding solutions. In our setting, the interval discrimination was performed between stimuli of different duration, rather than between waiting intervals as in previous studies [1,2,4]. These stimuli are not part of bees’ natural ecological environment. Therefore, bees’ ability to learn to discriminate between these artificial stimuli shows general learning abilities through the visual modality, rather than specialised foraging strategies.

In Experiment 1, two conditions were tested. The long cycle condition presented 1 vs 5 second stimuli (during a10 s cycle). The short cycle condition presented 0.5 vs 2.5 second stimuli (during a 5 s cycle). Each flashing stimulus was presented once per cycle. Bees learned to discriminate between the stimuli in both conditions. They learned both when the short and when the long stimulus was rewarded, showing that they were not driven only by spontaneous preferences and phototaxis [31,32]. Since each cycle in Experiment 1 presented the long stimulus five times longer than the short stimulus, to identify the position of the sucrose reward bees may have relied on interval duration difference or on the total amount of stimulation received. To check whether bees could solve the task based on duration differences only, we ran a second experiment in which the total amount of stimulation was matched for both stimuli, by having stimuli flash on and off throughout the cycle.

In Experiment 2, we used 0.5 vs 2.5 second stimuli, as in Experiment 1, but total amount of light per cycle light was matched between stimuli (2.5 seconds of light for both stimuli, Fig. 1C). Bees learned to discriminate between the stimuli, showing that they were using the duration as a cue to identify the position of the reward.

Previous research has shown that bumblebees (*Bombus impatiens*) can learn to expect a reward, measured by extension of the proboscis, after a given interval of 6, 12 or 36 seconds) [18]. Wasps can discriminate between durations of 5 and 30 minutes [17]. These studies have in common the demonstration of an ability to track the interval between or before rewards (the interval was used as conditioned stimulus). Other work has shown that bees are able to optimise foraging bouts by learning the rates at which nectar is replenished in flowers [1,2,4]. The present study shows that bees are also able to use time cues inherent to the predictor of reward (the conditioned stimulus itself).

The ability of insects to process interval duration originating from outside the organism has been studied naturalistic contexts, such as cricket calls [8]. While crickets have evolved to recognise calls for recognition of conspecifics and reproduction, there is no obvious ecological basis for interval discrimination of visual stimuli in bees. This points to a general learning ability in bees, that can extend to temporal processing to novel visual situations.

The ability of insects to discriminate between temporal stimuli outside their ecological niche suggests that interval duration may be a widespread phenomenon across animal species. It has been suggested that temporal computations might be an inherent property of neural circuits [33,34]. Whether or not general neural dynamics underly our results, bees’ ability to discriminate between interval durations in non-naturalistic stimuli point to domain-general skills.

Temporal cognition abilities might be linked to spatial skills. For instance, during flight bees use the rate of visual stimulation to measure speed and height [35]. Visual stimuli that intermittently appear in time might have similarities with the stimuli that flow across space during motion. A connection between the neural encoding of time and space has already been identified at the neuronal level in vertebrate species [36,37]. Future studies should explore the underlying basis of time encoding and its connection to space processing in insects.

Overall, our findings show that bumblebees can use visual processing to discriminate between durations of external stimuli, independent of reward rates. This suggests that temporal cognition in bees extends beyond foraging strategies and relies on domain-general mechanisms. The ability to track time in a non-naturalistic setting highlights a level of cognitive flexibility that warrants further investigation at the behavioural, computational and neural levels. Future research should explore whether similar mechanisms underpin time and space encoding in insects, as observed in vertebrates, and how these computations contribute to adaptive behavior across ecological and artificial contexts. By establishing robust methods for testing interval discrimination in bees, this study opens new avenues for understanding the fundamental principles of time perception in invertebrates.

## Acknowledgements

We thank Charlotte Lockwood for her excellent support with bee handling procedures.

## Data accessibility

All data are available in the main text or the electronic supplementary material.

## Declaration of AI use

We have not used AI-assisted technologies in creating this article.

## Authors’ contributions

A.D.: conceptualisation, investigation, formal analysis, visualisation, data curation, writing—original draft, writing—review and editing; I.N.: investigation, methodology, validation, writing—review and editing; A.O.L.M.; conceptualisation, investigation, methodology, validation, writing—review and editing; L.C.; conceptualisation, funding acquisition, supervision, writing—review and editing; E.V.: conceptualisation, funding acquisition, methodology, resources, supervision, writing—review and editing All authors gave final approval for publication and agreed to be held accountable for the work performed therein.

